# Development and Validation of a New Walking Pace Function Using Crowd-Sourced GPS Data

**DOI:** 10.1101/188417

**Authors:** Paul D.P. Pharoah

## Abstract

There are several functions that hikers can use to predict walking time based on elevation change or slope of the ground. The most commonly used is the Naismith function that was first published over 100 years ago. The availability of GPS devices to record tracks now make it possible to evaluate the performance of walking time functions. Four data sources were used: 98 tracks downloaded from the Wikiloc web site; 55 tracks recorded by the author; 19 tracks recorded by the blogger Iron Hiker; and 20 tracks recorded by the blogger Hiking Guy. The ․gpx files were processed to generate segements of ~100m in length, with the associated segment duration and elevation change. The association between walking pace and elevation change was assessed in the Wikiloc data using linear spline regression. The performance of the linear spline function was then compared with the Naismith, Tobler and Laingmuir functions. The linear spline performed the best, but all four performed reasonably well. While the linear spline function could easily be programmed into the software of standard GPS devices, the Naismith function provides a simple-to-use rule-of-thumb for estimating walking time for a typical hike in the mountains.

**Funding:** None

**Acknowledgements:** I thank Chris Hazzard and Keith Wilson for sharing their hiking records and for helpful comments on the manuscript.

**Disclosure of interests:** The authors have no interest to disclose

## Introduction

There are several widely used methods for estimating average walking pace as a function of distance travelled and change in elevation. The most well know of these are Naismith’s Rule and the Tobler Hiking Function. The Naismith function was only intended for predicting pace on an incline and stated that the a man in fair condition on an easy expedition should allow 1 hour for every 3 mi (5 km) forward, plus 1 hour for every 2000 ft (600 m) of ascent (Naismith). In the article Naismith does not explicitly state that the time taken is only to be calculated for the uphill parts of a walk, and, given that the rule was described in the context of routes that start and finish at the same elevation, it is reasonable to assume that he intended that rule was for the calculation of time for the complete route based on total ascent and distance. Converting his rule into SI units

> Total time = 12*distance (km) + 10*total ascent (m) /100
>
> Average pace (minutes per km) = 12 + 10*total ascent / (100 * distance (km))

The Naismith Rule implies equivalence of 3 miles and 2000ft climb (Scarf, 2008) and so the rule can be restated as

> Equivalent distance = total horizontal distance + 7.92*vertical distance ascended

Again, this formula would be expected to apply to a route with an equal amount of ascending and descending. Several corrections to the Naismith rule have been proposed. Tranter applied a fitness and fatigue function to the. Fitness is measurd by the time in minutes taken to climb 300m in a 800m linear walk with 15 being very fit and 50 being unfit. Tranter’s corrections also have further variations: walking with a headwind, a 20kg load and at night all mean coming down a fitness level, and terrain type can mean either one or two levels down on the scale.

However, the Naismith rule does not provide specific guidance on pace going uphill or pace going downhill and applying the rule to a route that starts and finishes at a different elevation makes calculation of expected time dependent on the underlying assumption about how the Naismith rule applies to pace on uphill and downgradients. It is often assumed that the Naismith time penalty only applied to uphill parts of a route and that downhill pace is the same as pace on level ground. Thus,

> Total uphill time = 12*distance on positive gradient + 10*total ascent (m) /100 +
>
> Total downhill time = 12*distance on negative gradient

This pace function is shown in **Supplementary Figure 1** and I will refer to this as the Naismith A function. Under this assumtion a 10km route with a net gain of elevation of 1000m would be expected to take 220 minutes wherea a route with a net loss of elevation of 1000m would be expected to take 120 minutes. On the other hand, or it could be assumed that pace was equally affetced by net gain and net loss of elevation so that one should allow for 30 minutes for every 2000 ft of ascent and 30 minutes for every 2000 ft of descent (Supplementary Figure 1). I will refer to this as the Naismith B function.

> Total uphill time = 12*distance on positive gradient + 5*total ascent (m) /100 +
>
> Total downhill time = 12*distance on negative gradient + 5*total descent (m) /100

Under this assumption a 10km route with either a net gain or a net loss of elevation of 1000m would be expected to take 170 minutes. The Naismith rule is also compatible with any number of intermediates between the A and B functions in which the pace is reduced by a down hill gradient, but to a lesser extent than an uphill gradient of the same slop.

Langmuir proposed an addition to the Naismith rule for downhill walking with base speed of 5 km/h applying for slopes of up to 5 degress and makes the following further refinements for going downhill. For slopes between5 and 12 degress the walker should subtract 10 minutes for every 300 meters of descent for slopes between 5 degrees and 12 degrees and then then add 10 minutes for every 300 meters of descent for slopes greater than 12 degrees (Supplemenatry Figure 1) (Langmuir, 1984).

The Tobler hiking function is an exponential function that also explicitly models pace according to whether the gradient is uphill or downhill according to the formula

> Speed (km per hour) = 6*exp {−3.5*abs(S + 0.05)}

where S is the slope of the climb. It predicts a maximum speed of 6 km per hour or 10 minutes per km on a down slope of 5 per cent.

Few published studies have reported on the accuracy of the predictions of various hill walking functions using empirical data. Scarf analysed data from record finishing times for British fell races to develop a model for record time as a function of total distance and total ascent. Like Naismith this approach does not explicitly model the ascent and the descent separately. He also discussed the Naismith function assuming that the pace at all downhill gradients is equal to that on the flat (Naismith A) and reported that fell running records provide supporting evidence for the Naismith function (Scarf, 2007).

More recently, Kay compared six functional forms for the shape of the association between gradient and pace for elite male hill runners using the record times from 91 uphill and 15 downhill races to obtain a function for elite male hill runners. These included functions of the form of the Tobler function and the Naismith A function. He also evaluated a piecewise linear model, which would include the Naismith B model and quadratic cubic and quartic functions. He did not compare the functions derived from the data with a direct application of the Naismith or Toble functions, but it is of note that the piecewise linear model fit the data the best (Kay, 2012).

There are no large scale analyses investigating the accuracy of the walking time predicted by different functions using empirical data for the “ordinary” walker. The aim of this study was to utilise GPS data recorded by walkers to evaluate the accuracy of walking time predicted by the Naismith rule, the Tobler function and to compare these with a function derived directly from the available GPS data.

## Data extraction and analysis

Four sources of data were used: i) a pseudo-random set of recordings downloaded from from www.wikiloc.com (Wikiloc), a website where individuals can uploading and share GPS tracks; ii) the authors own recording of 67 walks from around the world done since 2010; ii) recordings of 20 walks made by Iron Hiker (http://ironhiker.blogspot.com/) in California USA; and iv). Recordings of 20 walks made by Hiking Guy (https://hikingguy.com/) in Califronia USA. There are over two million tracks shared on the website. Wikiloc recordings were selected by first applying the filters: i) recorded on GPS device; ii) distance between 10 and 25 miles iii) total elevation gain over 1500 ft. Tracks were downloaded attempting to pick examples from around the world after excluding any tracks with obvious data signal problems based on visible linear segments from the track profile shown on the web page. Tracks were subsequently excluded if the average pace was less than 10 minutes per km suggestive of running, tracks recorded over a total time of greater than 16 hours, less than 5 km cumulative distance of eligible segments available for analysis (see below). This process was repeated until 102 representative tracks were downloaded.

### Data extraction

Latitude, longitude, elevation and time were extracted from each ․gpx file using the {plotKML}readGPX function in R. The distance between points was estimated using the pointDistance function which calculate the geographic distance between two (sets of) points on the World Geodetic System ellipsoid. Each track was then split into segments of approximately 100m in distance in which the points nearest to a cumulative distance of x metres were selected where x equals 100, 200, 300….final cumulative distance. The walking pace between points was calculated from the cumulative distance between the individuals points in the segment (close to 100m by definition) and the cumulative time between the points. This was converted to a pace of minutes per kilometre. The net elevation bewteen points was calculated as the elevation different between the first point of the 100m segment and the last point of the 100m segment.

Segments were excluded from further analysis if: Pace less than 8 minutes per km suggestive of jogging rather than walking; segment pace greater than 50 minutes per km as these are very likely to be due to a pause in walking during one or more consecutive segments; two consecutive segments where pace preater than 40 minutes per km; a change in absolute elevation of greater than 70 m.

### Data analysis

For each segment the expected duration based on the Naismith rule A, Naismith rule B, the Laingmuir function and the Tobler function were calculated from the segment distance and duration. The Wikiloc data set was used to derive a function for walking pace based on the GPS data.

A wide number of variables are likely to affect walking pace including characteristics of the terrian such as slope, altitude, prior distance walked, type of terrain, ambient temperature and humidity and wind speed, and characteristics of the individual such as age, load carried and individual fitness. However data with such detailed information are not easily available and so this analysis was restricted to the effect of slope, altitude and prior distance walked on typical walking pace. It is likely that the effects of distance, altitude and slope are confounded by each other. Therefore a multi-variable model was therefore built using the Wikiloc data in order to establish the independent effect of each of these variables. Segment pace (minutes per km) was the dependent variable with slope (change in elevation in metres over 100m segment), altitude, distance and data set the independent variables. Two approaches were used to model the relationship between pace and slope. First linear spline regression models with either a single knot or with three knots were compared. Five single knotmodels were evaluated with knots at elevation gains of −10m, −8m, −6m, −4m, −2m, 0m, 2m, 4m, 6m, 8m and 10m. The three knot models all had a knot at 0m with different combinations of knots for the down and up slopes. Because of the skewed distribution of the walking pace data quantile regression (median) rather than ordinary least squares regression was used. The second approach used multivariable fractional polynomials were used to model the associations of elevation change and altitude as these relationships may be non-linear. Distance was modelled as a factor variable in four groups (0-5km, 5−10km, 10−20km and 20+km).

The Author data set, the Iron Hiker data set and the Hiking Guy data set provided data for independent validation of the best fitting function derived using the Wikiloc data.

## Results

Twelve of the tracks from the author data set, one of the Iron Hiker tracks and three from the Wikiloc data set were excluded after applying the criteria described above. A total of 193 tracks comprising 28,517 segments of ~100m and 2,972 km of walking were included in the analysis. The average length of the walks was 15.4 km and the average ascent was 858m with 843m of descent. **Supplementary Figure 2** shows the location of each track by data set.

The association between walking pace and elevation change for each 100m segment by data set is shown in **Supplementary Figure 3**. This demonstrates a clear U-shaped relationship in all four data sets. The large number of outliers for elevation changes of between −20m and +20m are likely to represent segments in which the walker paused or rested. **Supplementary Figure 4** shows the association between walking pace and altitude for up slopes and down slopes by data set. Again, the pattern is consistent across all three data sets with an approximately linear reduction in walking pace for every 1000m of altitude. It might be expected that this relationship would differ between the segments on an up slope and segments on a down slope as the effects of oxygen deprivation might be expected to be greater when going uphill. However no clear difference is apparent. **Supplementary Supplementary** shows the relationship between walking pace and distance travelled. While it might be expected that pace would slow down as cumulative distance increases and fatigue sets in, there appears to be a slight increase in pace with distance for both the Wikiloc data and the author data.

The best fitting of the linear spline models was the model with knots at −10m, 0m and 10m elevation change over 100m. This model fit substantially better than the best fitting multivariable fractional polynomial model (log likelihood −41,260, 6df compared to −41,704, 5df). The coefficients from the linear spline model are shown in **Table 1**.

**Table 1:**
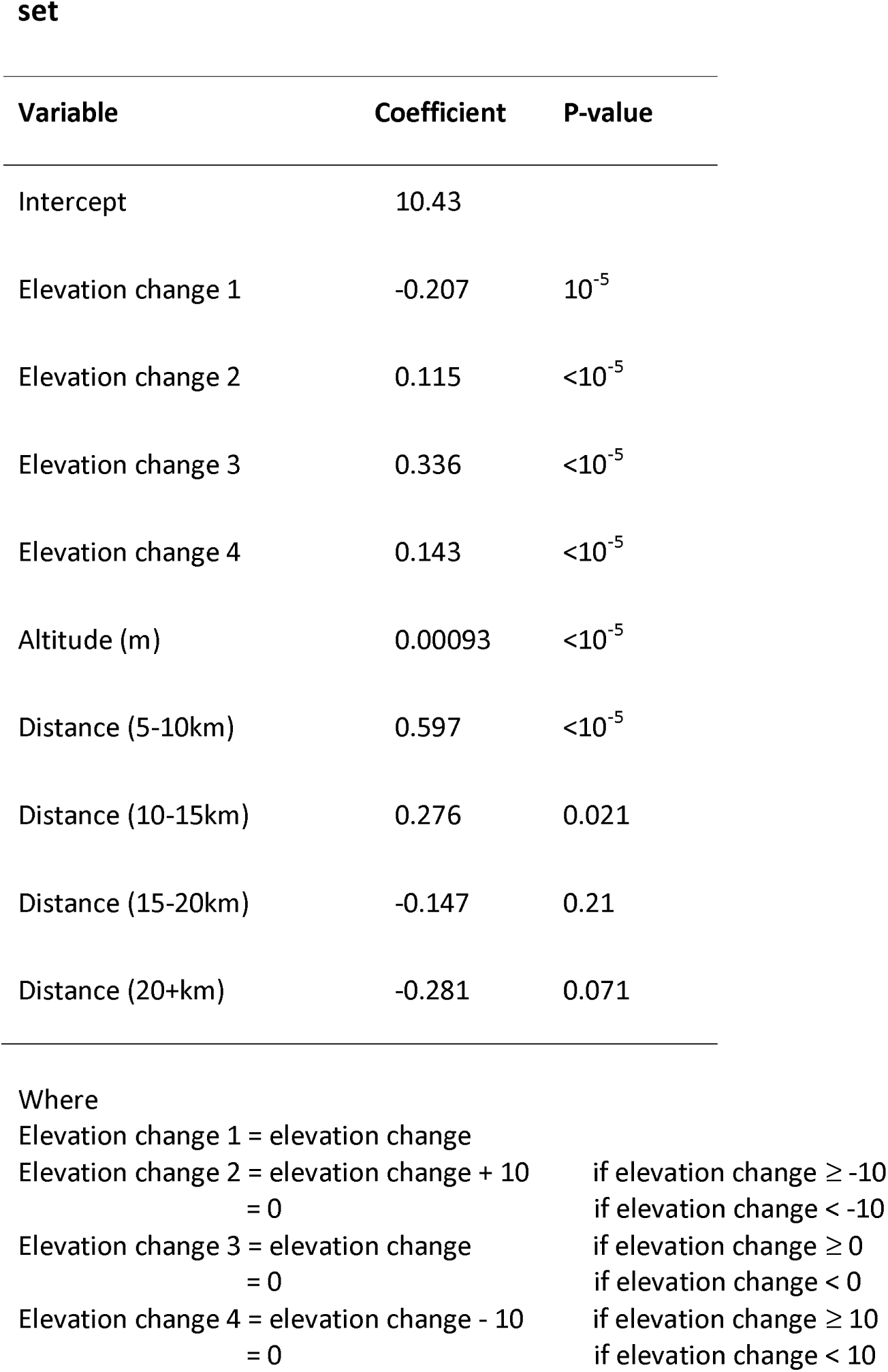
Coefficients from the best fitting linear spline model fit to the Wikiloc data set

Maximum pace is 12 minutes and 13 seconds per kilometer which occurs on level ground. Expected walking pace on an incline of 20 per cent (20m gain over 100m) at sea level is 19 minutes 42 seconds per km and this increases to 22 minutes 30 seconds at an altitude of 3000m. Walking pace on a decline of 20 per cent at sea level is predicted to be 15 minutes 13 seconds per km. **Figure 1** is a scatter plot of walking pace against altitude for each of the four data sets with the line of best fit derived from the Wikiloc data and the Naismith B function superimposed.

**Figure 1:**
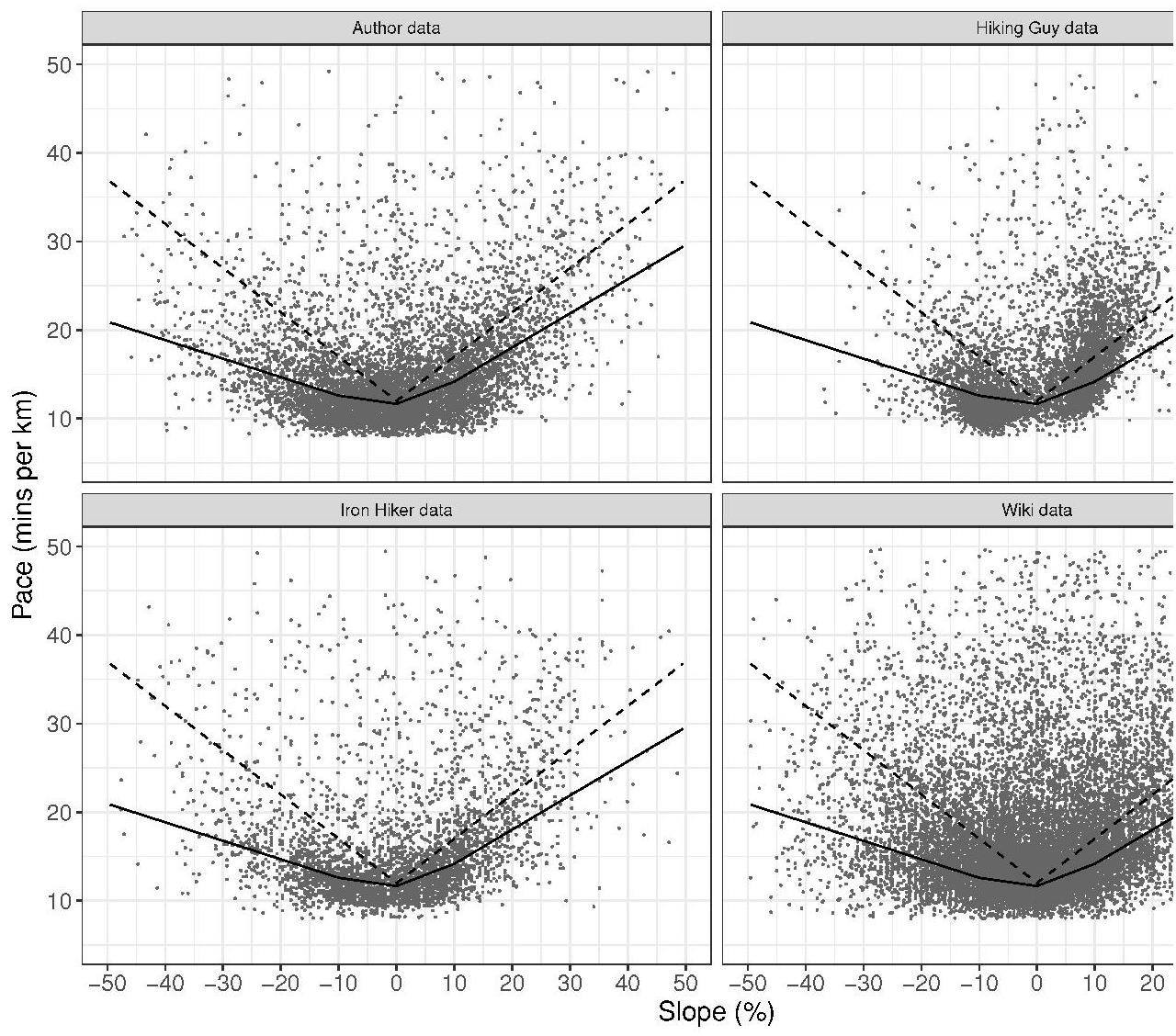
Scatterplot of walking pace for each 100m segment by elevation change for each data set. Superimposed lines are the Naismith Rule B function (dashed) and the best fitting linear spline based on the Wikiloc data (solid).

One of the major problems with the analysis of GPS data is the variation inherent in each data point recorded in a track. Each ~100m chunk is made up of an average of 5 recorded segments. At each of the six recording points marking the start and the end of the chunk there will be errors in the GPS location and in the conversion of that location into distance and elevation change. In addition, a substantial portion of the variation in pace between chunks will simply be due to the walker pausing - a 2 minute pause in the middle of a 100m chunk could change the walking pace from 15 minutes per km to 35 minutes per km for example. It is likely that a large number of the chunks with pace slower than 30 minutes per km were associated with a walking pause. Of course, on steeper inclines or a high altitude, a pause may simply be to get one’s breath back.

However, the errors from point-to-point in a recording, and for each multi-segment chunk are likely to even out over several kilometres of walking. **Table 2** shows the overall fit of the different functions for the total walking time for the model development data and the model validation data and **Figure 2** shows the the actual walking time for each of the 193 walks plotted against against the predicted walking time for the best-fit linear spline model model fitted to the Wikiloc data, the Naismith B function, theTobler function and the Laingmuir function.

**Table 2:** Comparison of predicted and observed walking times for four prediction functions by data set

| Function | Model development data |  | Model validation data |  |
| --- | --- | --- | --- | --- |
|  | Predicted - observed | (Predicted – observed)<sup>2</sup> | Predicted - observed | (Predicted – observed)<sup>2</sup> |
| Total |  |  |  |  |
| Observed | 24,382 |  | 21940 |  |
| Linear spline | -209 | 102,012 | 440 | 78,216 |
| Naismith B | 2213 | 155,220 | 3539 | 243,404 |
| Tobler | 567 | 145,509 | 597 | 99,473 |
| Laingmuir | 2128 | 215,485 | 2887 | 181,793 |
| Up segments |  |  |  |  |
| Observed | 12297 |  | 12009 |  |
| Linear spline | 102 | 28,025 | -302 | 39,065 |
| Naismith B | 644 | 29,023 | 981 | 44,682 |
| Tobler | 1703 | 79,261 | 1094 | 56,351 |
| Laingmuir | 4962 | 361,220 | 4448 | 289,256 |
| Down segments |  |  |  |  |
| Observed | 12086 |  | 9932 |  |
| Linear spline | -311 | 31,215 | 742 | 34,159 |
| Naismith B | 1569 | 63,606 | 2558 | 114,578 |
| Tobler | -1136 | 47,770 | -496 | 19,392 |
| Laingmuir | -2834 | 120,098 | -1561 | 51,994 |

**Figure 2:**
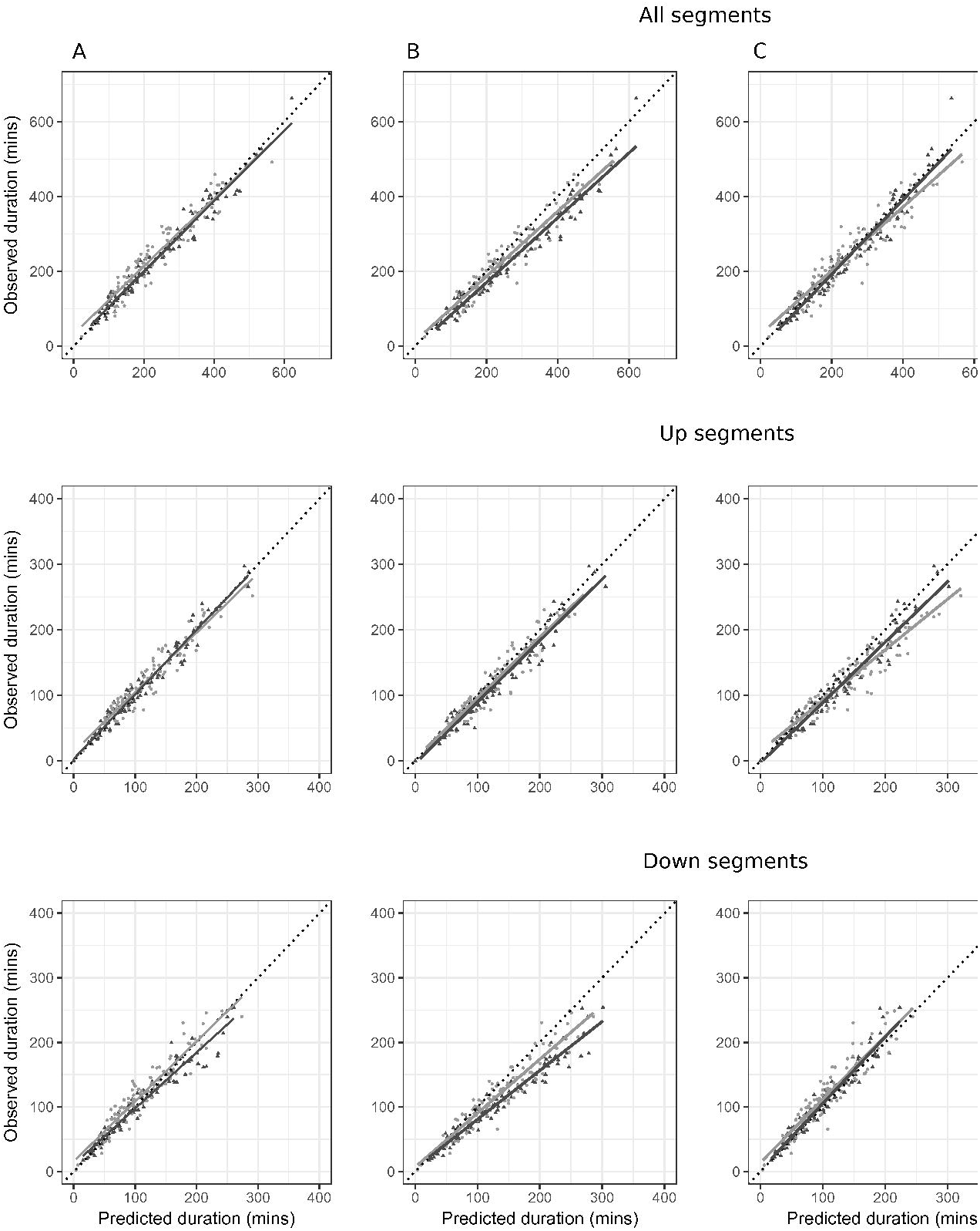
Observed duration (total, up segments and down segments) in minutes for 193 tracks compared to predicted duration for (A) best fit linear spline model (B) Naismith B function (C) Tobler function and (D) Laingmuir function. The dark grey line shows the line of best fit for the model development (Wikiloc) data and the light geey line shows the line of best fit for the validation data. The dotted line shows the expected line for a perfect model (observed = predicted).

Based on the sum of the squared difference between predicted and actual walking times as a measure of fit, the linear spline function performed best on the model development data set for total walking time, and for walking time on the up and dwon segments. The linear spline model also performed best for total walking time and up segment walking time in the model validation data, with the Tobler function being the best performing model on the dwon segments. On average the linear spline model resulted in a small overstimation of total walking time in the validation data (22,380 minutes predicted v 21,940 minutes observed), with a slight underestimation of walking time on the down segments and an overestimation on the up segments.

## Discussion

The analyses presented here have used publicly available GPS tracking data to evaluate two key factors that influence walking pace - slope and altitude. The data used for this analysis are likely to represent a wide range of different types of terrain for hikes undertaken in a wide variety of conditions at different times of the year. The data are also likely to represent the walking pace of the hiking enthusiast rather than the casual walker given that it is the enthusiast who is likely to keep a record of their walks and upload them onto a public web site.

Limited annotation of the data prevented the evaluation of several other factors that are likely to be important. For example, terrain and conditions underfoot are likely to be important. Every walker knows that meterological factors such as ambient temperature, wind, rain and snow can all affect pace. Another factor which obviously contributes to walking time is the age and fitness of the hillwalker. Tranter’s correction to the Naismith function takes an individual’s fitness into account. The Tranter correction is in the form of a table, where the basic Naismith estimate for a route is modified by a factor which is dependent on individual fitness level; this in turn can be determined by recording the time taken to climb a set height (300m) over a set distance (800m) at normal walking space. Tranter’s correction also takes into account the effects of fatigue with a reduction in pace for longer excursions, although the data in this paper suggests that fatigue is not a major factor.

In summary, the analysis presented here has shown that a simple linear spline regression model fit using publicly available GPS tracking data outperforms the most commonly used walking time function in an independent validation data set. However, while the model is conceptually simple it does not convert into a simple rule-of-thumb that could be easily applied to predict total walking time for an excursion. Nevertheless, given the popularity of GPS navigation devices it would be comparatively straightforward to programme the predicted walking time for a planned excursion or for any given segment of an excursion. The Tobler and Laingmuir functions are equally difficult to apply and perform less well than the linear spline model. They would appear to have little to recommend them for normal usage. On the other hand the Naismith function performs reasonably well at predicting total walking duration based on total elevation gained. For walks that start and end at the same point either the Naismith A or the Naismith B function can be used. If, the walk is a point-to-point or a sement of a round trip, with a difference in total elevation gained and total elevation lost, the Naismith B function is a more reliable option.

## List of Supplementary figures

**Supplementary Figure 1:** Walking pace (minutes per km) by slope (elevation change in metres over 100m) for the Naismith A function, Naismith B function, Langmuir function (Naithmith A with negative gradient correction) and Tobler function.

**Supplementary Figure 2:** Location of tracks used in the analysis. Black points model development data (Wikiloc), grey points model validation data (author, Iron Hiker and Hiking Guy).

**Supplementary Figure 3:** Boxplot of walking pace by elevation change for each 100m segment and data set

**Supplementary Figure 4:** Boxplot of walking pace by altitude for each 100m segment and data set

**Supplementary Figure 5:** Boxplot of walking pace by distance walked for each 100m segment and data set. Up sloping and downsloping segments shown separately

